# Single-molecule sizing through nano-cavity confinement

**DOI:** 10.1101/2021.12.13.472369

**Authors:** Raphaël P. B. Jacquat, Georg Krainer, Quentin A. E. Peter, Ali Nawaz Babar, Oliver Vanderpoorten, Catherine K. Xu, Timothy J. Welsh, Clemens F. Kaminski, Ulrich F. Keyser, Jeremy J. Baumberg, Tuomas P. J. Knowles

## Abstract

An approach relying on nano-cavity confinement is developed in this paper for the sizing of nanoscale particles and single biomolecules in solution. The approach, termed nano-cavity diffusional sizing (NDS), measures particle residence times within fluidic nano-cavities to determine their hydrodynamic radii. Using theoretical modeling and simulation, we show that the residence time of particles within nano-cavities above a critical timescale depends on the diffusion coefficient of the particle, which allows estimation of the particle’s size. We demonstrate this approach experimentally through measurement of particle residence times within nano-fluidic cavities using single-molecule confocal microscopy. Our data show that the residence times scale linearly with the sizes of nanoscale colloids, protein aggregates and single DNA oligonucleotides. NDS thus constitutes a new single molecule optofluidic approach that allows rapid and quantitative sizing of nanoscale particles for potential application in nanobiotechnology, biophysics, and clinical diagnostics.

**Table of content graphic:** 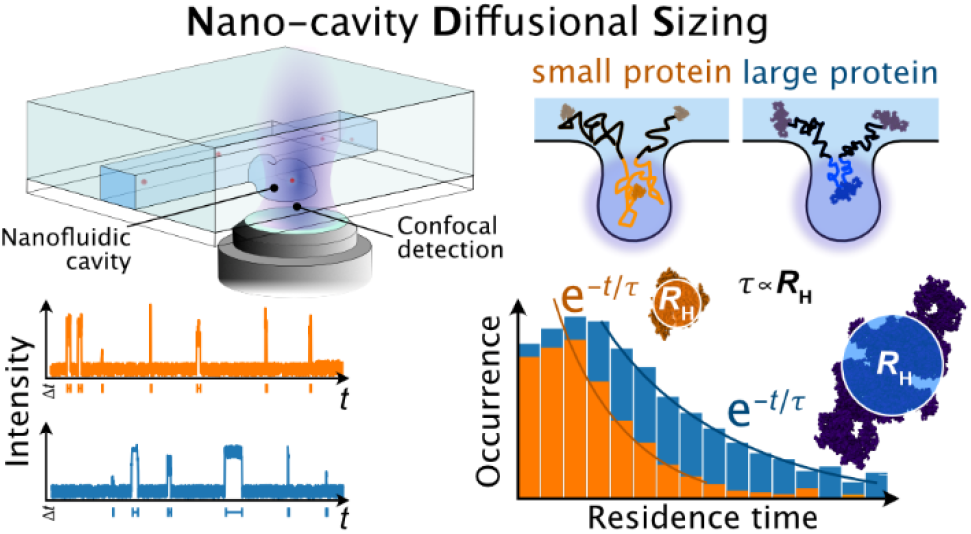

---

Many important biomolecules, including proteins and protein assemblies, as well as natural and synthetic biopolymers and colloids, have sizes in the nanometer range.^1,2^ Achieving rapid, accurate, and reliable measurement of their sizes under native solution conditions has therefore become a key objective in many areas of current research including nanobiotechnology, biophysics, and clinical diagnostics.^3–5^ For example, sizing of proteinaceous particles at nanometer scales is critical in studies that further our understanding of protein misfolding and aggregation processes, which lie at the heart of a wide range of human diseases.^6–8^ Moreover, characterizing the assembly state of biomacromolecules is important when assessing, for example, biopharmaceutical product stability and efficacy of proteins or biocolloids in drug delivery systems and formulations.^9–11^ Sizing techniques are therefore considered ‘workhorse’ methods in many areas of fundamental and applied science.^3,12^ Hence, the development of experimental approaches for high sensitivity detection and characterization of nanoscale entities in the fluid phase remains an area of great current interest.

Several techniques are available in order to measure the nanoscale size of proteins and nano-colloids in solution.^3–5^ Most of them are based on determining the particle’s diffusion coefficient *D* in solution, which is related via the Stokes-Einstein equation to the hydrodynamic radius *R*_H_ of the particle. One of the most widely used techniques is dynamic light scattering (DLS).^13^ Other widely used methods include nuclear magnetic resonance (NMR)-based techniques (e.g., pulse-field gradient NMR),^14^ chromatographic techniques,^15^ and surface deposition microscopy, like atomic force microscopy (AFM) or scanning/transmission electron microscopy (SEM/TEM).^16,17^ These techniques suffer from relatively high sample consumption and long acquisitions times or surface immobilization, and often require sophisticated instrumentation.

In recent years, a number of techniques have been established that operate with minimal sample requirements and sensitivities down to the single molecule regime, and operate directly in solution. These include microfluidic techniques^18,19^ such as microfluidic diffusional sizing (MDS),^8,20^ Taylor dispersion analysis (TDA),^21^ nanoparticle tracking analysis (NTA)^22^ and single molecule techniques such as fluorescence correlation spectroscopy (FCS)^23^ or a combination of interferometric scattering (iSCAT) microscopy^24^ with electrostatic trapping.^25^ Such fluidic and single molecule-based approaches offer great potential for the sizing of nanoparticles and nano-colloids in solution, however, they are often limited in the size range that can be detected. In fact, there is a critical analytical gap in the size range that can be detected by such methods. NTA and other SMT/SPT-based techniques perform robustly only for particle sizes above approximately 50 nm in radius, while MDS and FCS are mostly limited to particle radii below 10–20 nm. Moreover, techniques such as MDS, TDS, and FCS require complex models to analyze the size distributions. Thus, approaches that can measure a wide range of particle sizes in solution with minimal sample consumption within the range of a few nanometers up to the hundreds of nanometer size range are much sought after.

Recently, we have reported the fabrication of nanofluidic devices with nano-cavity confinement functionalities that enable single-molecule studies at prolonged observational-time scales to analyze and detect nanoparticles and protein assemblies in solution without the need for surface immobilisation.^26^ Using confocal fluorescence burst detection, we therein made the striking observation that various nanoscale-sized particles exhibited a marked increase in average residence times in the detection volume of the nano-cavities and that this residence time increase scales with the size of the particles. Here, we take advantage of these observations to develop a quantitative single-molecule approach for the size determination of nanoscale particles in solution.

The approach introduced here, termed nano-cavity diffusional sizing (NDS), is based on the idea that the sizes of nanoparticles can be extracted from measurements of particle residence times within fluidic nano-cavities. Through theoretical modeling and simulation, we show that the residence time of particles within nano-cavities above a critical timescale depends on the diffusion coefficient of the particle, which allows estimation of the particle’s size. We experimentally demonstrate the approach through measurement of particle residence time distributions within nano-fluidic cavities using single-molecule confocal microscopy.

The main concept and experimental realization of the NDS approach is outlined in **Figure 1**. The observation volume of a confocal microscope is placed within one of the trapping cavities of a nanofluidic device, which is filled with aqueous solution containing the biomolecule or colloid of interest (**Figure 1a**). The chip design is shown in **Figure 1b**. To extract the sizes of single molecules, time trajectories of a large number of single particles diffusing into and out of the observation volume are recorded, and residence time distributions are extracted (**Figure 1c**). Fitting of the obtained residence time histograms with a quantitative model provides *R*_H_ and, thus, the size of the nanoparticle of interest (**Figure 1c**).

**Figure 1.**
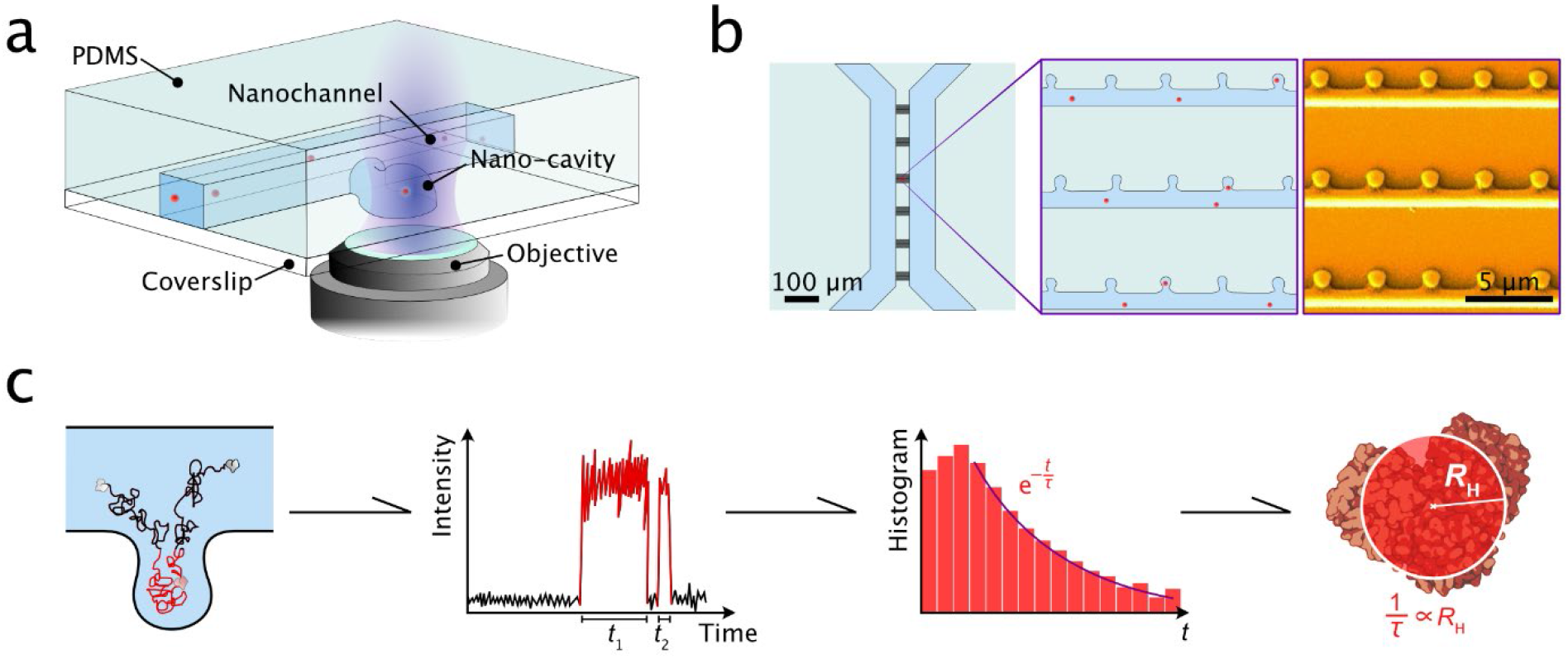
Principle of nano-cavity diffusional sizing (NDS). **(a)** Schematic illustration of the experimental implementation of NDS approach. The detection volume of a confocal microscope is placed inside the nano-cavity of the nanofluidic chip and residence times of single particles are recorded as they diffuse in and out of the nano-cavity. The nanofluidic chip is fabricated by hybrid lithography (see Methods). Single particles are shown in red. **(b)** Schematic of the nanofluidic chip used for NDS measurements. The nano-cavities are located adjacent to nanofluidic channels on a microfluidic chip. An SEM image of the chip is depicted in the right panel. Adapted with permission from Vanderpoorten et al.^26^. Copyright 2022 ACS. **(c)** Workflow of the sizing experiment. First, the particles are detected by confocal microscopy as they diffuse into and out of the nano-cavity (shown is a 2D representation of the nano-cavity with an adjacent nano-channel, as depicted in panel b). Then, the residence times *t* are extracted from the recorded time trace and binned in a residence time histogram. This histogram is then fit with an exponential function of the type 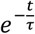, from which the hydrodynamic radius *R*_H_ can be extracted. The coefficient *τ* is inversely proportional to *R*_H_.

## Theory and Simulation

We first developed an analytical model to examine the diffusive behavior of particles within nano-cavities and to quantitatively describe how particle size relates to residence time. This allowed us to assess the scaling behavior of particle residence times and provided us with a theoretical framework for the analysis of our experimental results. We modeled the diffusion of particles as Brownian motion with reflective boundary conditions on the walls. No other potential was considered as we work under conditions where the direct electrostatic interactions between analyte particle and the cavity walls are negligeable because the particle is typically separated by a distance greater than the screening length. The nano-cavity, as shown in **Figure 1a**, was modeled as a cavity of rectangular shape (see 2D projection in **Figure 2a**), and is perpendicularly connected to adjacent nanochannels. The particle, in the model, can therefore only enter and exit the cavity by diffusing perpendicular to the nanochannel axis. Assuming that diffusion is isotropic (decorrelation in XYZ direction), the model, for the case of a rectangular cavity, can therefore be reduced to a one-dimensional (1D) diffusion problem.

**Figure 2.**
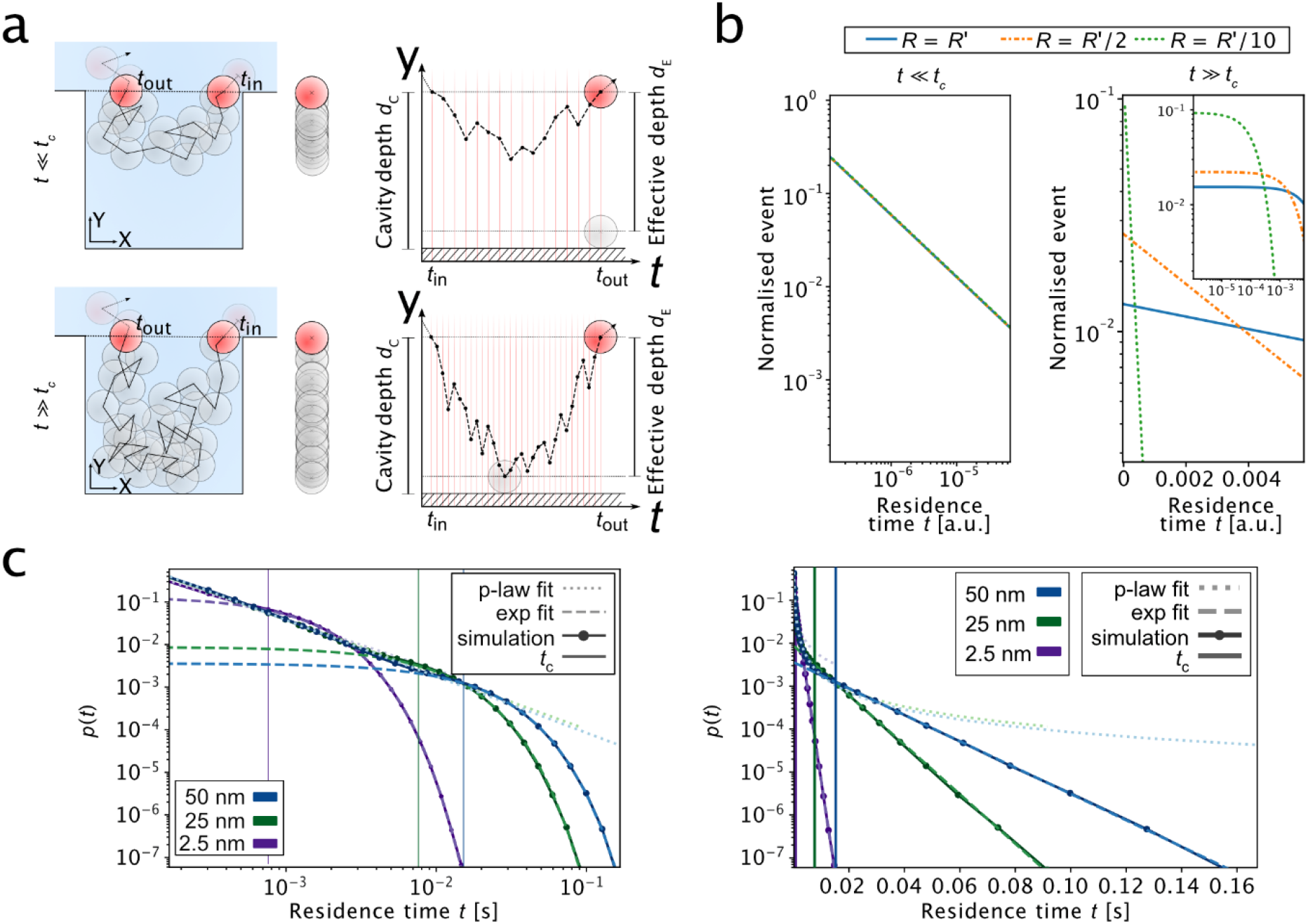
Theory and simulation of diffusion under nano-confinement. **(a)** 2D schematic representations of the nano-cavity model with particles entering and exiting the cavity (left), its corresponding 1D positions of particles along the Y-axis (centre panel) and its time trajectories along the Y-axis (right panels). For a particle diffusing within a nano-cavity, two diffusive scenarios can be distinguished: (i) the particle enters the cavity and exits it without reaching the bottom wall (top panels) or (ii) the particle enters the cavity and reaches the bottom of the well before exiting it (bottom panels). The right panels show displacement in time along the y-axis of the diffusion processes. The particle enters at time *t*_in_ and exits at time *t*_out_. The cavity depth is *d*_c_, and *d*_E_ denotes the effective cavity depth (*d*_E_ = *d*_c_ – *R*), with *R* being the radius of the particle. **(b)** Analytical modeling of the particle diffusion at short timescales (*t* << *t*_c_, left panel) and long timescales (*t* >> *t*_c_, right panel). Shown are residence time probability plots; *t*_c_ denotes the critical time (see main text). Shown are residence time probability plots. At short timescales (*t* << *t*_c_), the residence time is independent of the size of the particle. At long timescales (*t* >> *t*_c_), the residence time follows an exponential decay which depends on the particle’s size. Modeled were three particles with different radii *R*’ fixed at: *R* = *R*’ (blue), *R* = *R*’/2 (orange), and *R* = *R*’/10 (green). Diffusion was modeled as a 1D random walk. **(c)** Simulation results for the diffusion of particles within a nano-cavity. Shown are residence time probability plots (log–log plot, left panel; linear–log plot, right panel) for particles of different seizes (50 nm, blue; 25 nm, green; 2.5 nm, purple). At short residence time, particles are scale invariant. At long timescales, particle residence times exhibit an exponential decay, which is dependent on the size of the particle. Data points represent simulation results. Long and short dashed lines depict fits of the simulation data by power law and exponential functions, respectively.

The residence time of a particle is determined by the probability of the particle exiting the cavity over a given period of time. Representative time trajectories of a particle entering and exiting the cavity are shown in **Figure 2a**. The particle enters the cavity at time *t* = 0 and the time points to describe particle entry and exit are denoted as *t*_in_ and *t*_out_, which yields the residence time *t* = *t*_out_ – *t*_in_. The length of the cavity is *d*_*E*_, where *d*_*E*_ is the effective depth of the cavity and is given by the depth of the cavity *d*_*C*_ minus the particle radius *RR*:

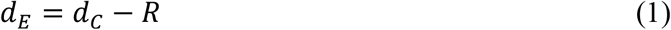

As depicted in **Figure 2a,b**, two regimes can be observed in the residence time distribution: a short and a long timescale regime. These two regimes are separated by a critical time *t*_c_, which corresponds to the mean time for a particle with diffusion coefficient *D* to reach the bottom wall of the nano-cavity according to:

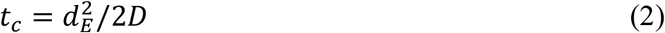

For short timescales, the particle resides within the cavity only for a short period of time such that it typically does not diffuse to the far end of the cavity (**Figure 2a, upper panels**). The probability distribution of the particle position inside the cavity is therefore concentrated near the original position. The distribution in this regime can be described by an unconstrained random walk model and is given by the first passage time density:

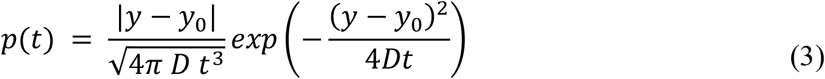

where *y* is the coordinate position between the inside/outside of the cavity (position where *t* = 0) and *y*_0_ is the distance that the particle diffuses inside the cavity in y-direction. For very small *Δy*, such that 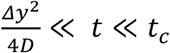, the density can be expressed as:

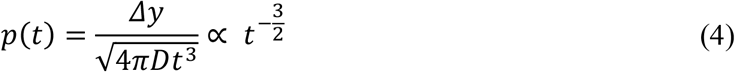

Accordingly, for the system at 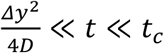, the residence time follows a power law, which is independent of *D*, because *Δy* scales with 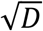 (see Supporting Information). This behavior is displayed in **Figure 2b, left panel** for particles with different diffusion coefficients.

For long timescales, the cavity space starts playing a role in the diffusion process, as the particle has enough time to explore the confined volume through diffusion. The free random walk model can therefore no longer be applied. As described in the Supporting Information and detailed in an analogous situation in Ref.^27^ (Equation 5.47 therein), the residence time distribution in this regime has an exponential dependence:

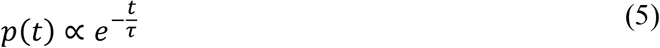

with the decay time being:

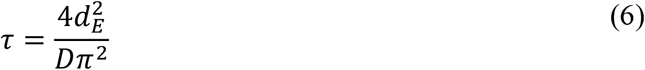

Accordingly, *τ* is inversely proportional to *D*. Hence, for the system at *t* ≫ *t*_*c*_ the size of the particle is linked to the decay time. This scaling behavior is shown in **Figure 2b, right panel** for particles with different diffusion coefficients. Residence time measurements at large timescales (*t* ≫ *t*_*c*_) allow estimations of sizes. This model can therefore size particles without the requirement to have an energetic contribution from any kind of a trapping free energy potential.

To corroborate our results from analytical modeling, we further performed numerical simulations of a particle 1D random walk within a nano-cavity to extract residence time probability distributions for differently sized particles. The simulations were performed using a reflective boundary condition for the wall on the bottom of the cavity. A Gaussian random number generator was used to simulate diffusive steps. Details of the simulations are given in the Methods section. Obtained residence time probability distributions are shown in **Figure 2c**. The results are consistent with the theory above, in that, at short timescales, the particle’s residence times follow a power law behavior 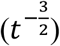, as evident in a linear decay in the log–log plot, whereas at long timescales, the residence time decays exponentially 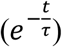, as evident in the linear decay in the linear–log plot. Fitting of the simulation results recovered the initial input values with a relative error of 0.5%, demonstrating the robustness of our analysis approach.

## Experimental demonstration

After having explored the possibility to size particles in nano-cavities on a theoretical basis, we next set out to demonstrate the NDS approach experimentally. Conceptually, the experimental implementation of the NDS approach involves the following steps: First, the durations of a large number of trapping events within nano-cavities is recorded, from which residence time decay histograms, which plot the occurrence of detected single particle residence times as a function of residence times, are generated. Then, by fitting this distribution with an exponential function, Eqs. 5 and 6 can be used to calculate the diffusion coefficient from the decay time of this exponential.

Based on these considerations, we set out to experimentally implement the NDS approach for the sizing of single particles in solution. We made use of a nano-fluidic device, previously developed in our laboratory, to measure particle residence times within nano-cavities.^26,28^ A schematic of the fluidic platform is shown in **Figure 3a, upper panel**. The device consists of arrays of nano-cavities which are connected to nanofluidic channels. These nanofluidic functionalities lie in between two microfluidic reservoirs with inlets and outlets that serve as fill ports for the sample solution. The nano-cavities are of cylindrical shape and have a radius of 350 nm and a height of 650 nm. The connecting nano-fluidic channels are 650 nm wide and 750 nm high. SEM images of the channel and nanocavity geometries are shown in **Figure 3a, lower panels**. The fabrication of the polydimethylsiloxane (PDMS)–silica devices by UV and 2-photon hybrid lithography (2PL) is detailed in the Methods section. For the detection of single particles, we made use of confocal fluorescence spectroscopy. Samples were excited with a continuous wave diode laser and their fluorescence was collected using avalanche photodiodes, which allowed us to readout the fluorescent signal of molecules with high sensitivity and monitor their residence times within nano-cavities with high temporal resolution. A schematic of the confocal microscope equipped with a motorized stage for precise placement of the confocal volume is shown in the Supporting Information.

**Figure 3.**
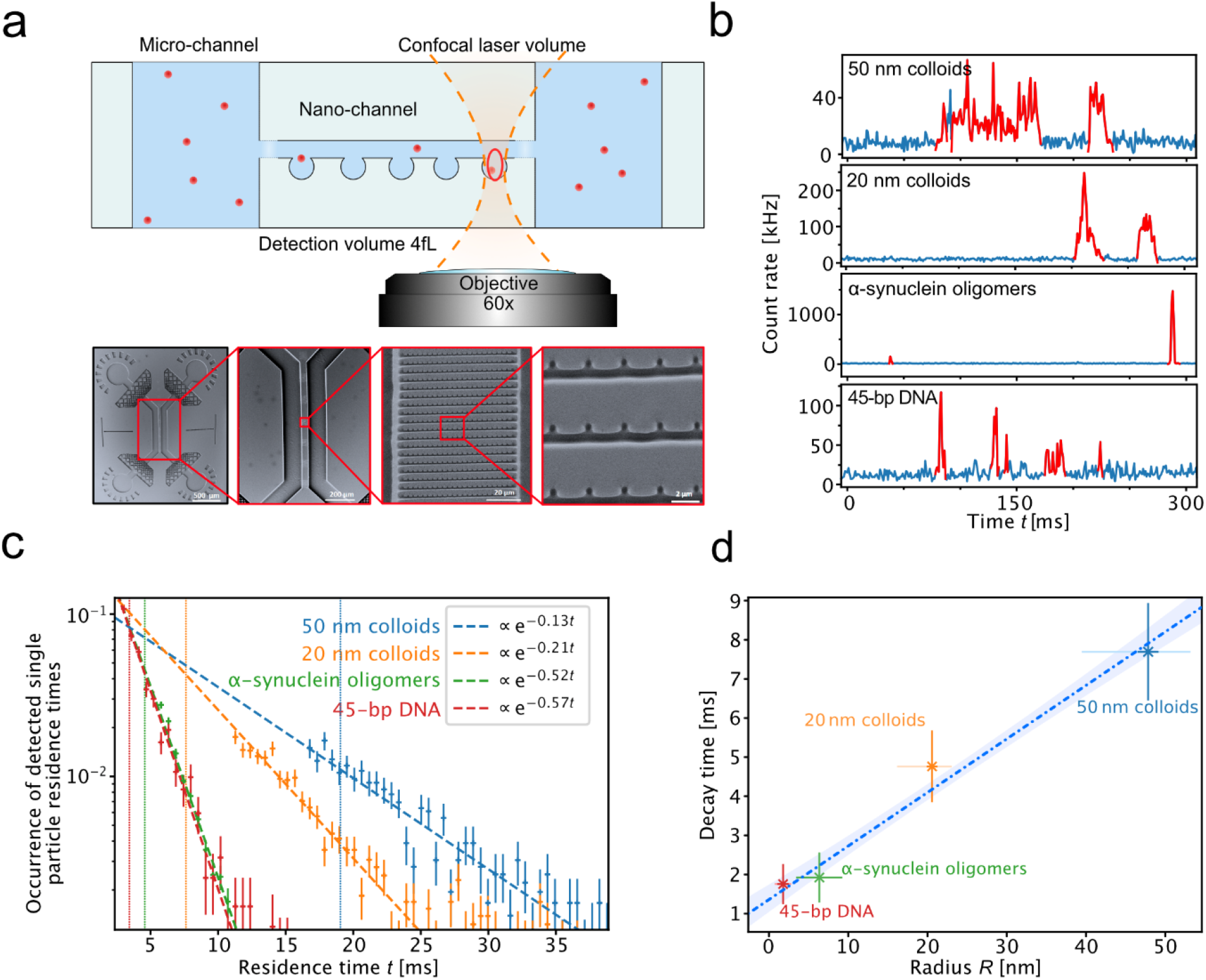
Nano-fluidic diffusional sizing (NDS) of single particles in solution. **(a)** Experimental setup of the NDS experiment. The observation volume of the confocal microscope is placed within a nano-cavity. Fluorescence of particles or biomolecules of interest are observed as they diffuse in and out of the confocal volume. Lower panel: SEM micrographs of the nanofluidic device with nano-cavity functionalities used in NDS experiments. SEM micrographs were adapted from Vanderpoorten et al.^26^. Copyright 2022 ACS. **(b)** Examples of time traces from single-molecule detection of nano-colloids, α-synuclein oligomers, and a DNA oligonucleotide within nano-cavities. Highlighted in red are the times when a molecule or particle was present within the confocal detection volume. The bin time is 1 ms in all traces. **(c)** Residence time decay histograms. The data were fit with an exponential function of the form 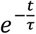. The slope of the curves gives the decay time. The dotted line corresponds to the critical time *t*_*c*_ for each species. The error bars are standard deviations of the Poisson distribution 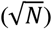, calculated from the number of events per bin (*N*). The following number of single molecule events were probed in order to create residence time histograms: 45-bp DNA: 969 events, α-synuclein oligomers: 1410 events, 50 nm colloids: 760 events, 20 nm colloids: 1137 events. **(d)** Extracted decay times versus hydrodynamic radii. The dotted line is a linear fit of the data points. Hydrodynamic radii for the colloids and the DNA were measured by dynamic light scattering (DLS) and for the oligomers by analytical ultracentrifugation (AUC). For a description of the error bars see Table 1.

Using this optofluidic platform, we probed the residence times of nanoscale particles for size determination by NDS. We performed measurements on fluorescent nanoscale colloids (50 and 20 nm in radius), fluorescently labeled oligomeric aggregates of the protein α-synuclein, and a fluorescently labeled DNA oligonucleotide (45 bp) (see Methods). See **Table 1** for details. Sample solutions were injected into the fluidic device and the confocal observation volume placed in the center of one of the nano-cavities of the device. After a short equilibration period to ensure hydrostatic balance, single molecule fluorescence of particles diffusing into and out of the nano-cavity were recorded. Exemplary time traces are shown in **Figure 3b**. As anticipated from our theoretical consideration, larger particles/molecules resided longer within the nano-cavities as compared to smaller ones. We measured hundreds to thousands of single molecule events (see **Figure 3c**). For each detected event, we extracted the associated residence times. Individual residence times were pooled in a histogram to obtain residence time histograms. Notably, due to the nature of the measurement, short residence events are under-sampled, which would create an artefact in the distribution (see Supporting Information). An adaptable threshold was therefore applied in order to represent residence times only at longer timescales. Moreover, residence times at short timescales are scale invariant; hence for size determination this regime can be omitted (see Theory and Simulations above).

**Table 1.**
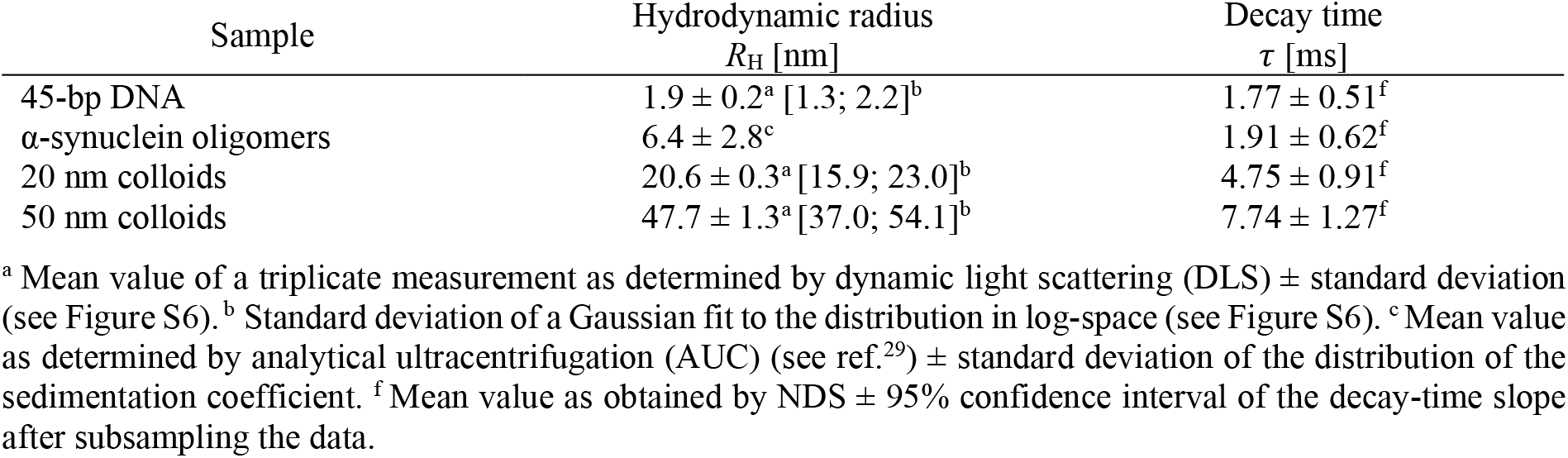
Samples used for NDS measurements.

| Sample | Hydrodynamic radius | Decay time |
| --- | --- | --- |
| | $R_H$ [nm] | $\tau$ [ms] |
| 45-bp DNA | $1.9 \pm 0.2^a$ [1.3; 2.2]<sup>b</sup> | $1.77 \pm 0.51^f$ |
| α-synuclein oligomers | $6.4 \pm 2.8^c$ | $1.91 \pm 0.62^f$ |
| 20 nm colloids | $20.6 \pm 0.3^a$ [15.9; 23.0]<sup>b</sup> | $4.75 \pm 0.91^f$ |
| 50 nm colloids | $47.7 \pm 1.3^a$ [37.0; 54.1]<sup>b</sup> | $7.74 \pm 1.27^f$ |

<sup>a</sup> Mean value of a triplicate measurement as determined by dynamic light scattering (DLS) ± standard deviation (see Figure S6). <sup>b</sup> Standard deviation of a Gaussian fit to the distribution in log-space (see Figure S6). <sup>c</sup> Mean value as determined by analytical ultracentrifugation (AUC) (see ref.<sup>29</sup>) ± standard deviation of the distribution of the sedimentation coefficient. <sup>f</sup> Mean value as obtained by NDS ± 95% confidence interval of the decay-time slope after subsampling the data.

The experimentally obtained residence time histograms for the four tested species are shown in **Figure 3c**. The residence time decays follow a linear behavior in the linear–log plot, as expected for an exponential behavior due to the biased random walk of the particles within the nano-cavity, as predicted from our theoretical modeling and simulations (see above).

Accordingly, we fitted the data with an exponential function of the form 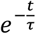. This allows extracting a decay time *τ*, which is proportional to the size of the particle according to the theory derived above, considering also the effective well depth. A plot of the extracted decay times versus the size of the particles yields a linear relation (**Figure 3d**), as anticipated from the theory. This linear relationship exemplifies the possibility to size particles in solution by NDS. Notably, while the theory above shows that the size of a particle can be directly recovered using a decay time histogram, in practice a calibration curve with size standards, as the one shown in **Figure 3d**, is required to correlate the decay times with particle size.

As exemplified, while NDS is able to measure particles from a few nanometers up to hundreds of nanometers, the method has its limitations. The largest size that can be measured with NDS is given by the size of the cavity. In the current configuration, the exclusion size of the cavity is on the order of 400 nm. Taking hydrodynamic coupling effects with the wall into account (see below), accurate sizing of particles with *R*_h_-values up this limit should be possible, and a different device design with a larger cavity volume should allow for the sizing of even bigger species. The smallest size that can be measured is determined by the minimal number of data points that would allow for a reliable extraction of the slope in the residence time decay histogram. While in theory there is no lower boundary—given that there is enough time to acquire data points in the decay histogram—in practice, the smallest species that can be sized will be determined by the experimental noise, especially at short decay times. The smallest species, measured in our experiments, was a 45-bp DNA sample with an *R*_h_-value of 1.9 nm, which exhibited a decay time of 1.77 ms and was determined from 15 data points (see **Figure 3c**). Assuming that at least 5 data points are necessary for a reliable determination, decay times much smaller than 1.5 ms, equivalent to a particle size of 1.13 nm, would become difficult to size reliably. Notably, the noise can be reduced by increasing the measurement time and hence the lower limit can be further improved potentially allowing for the sizing of sub-1-nm particles.

We further note that the nano-cavities created are not perfectly rectangular and their bottleneck-like structure at the entry/exit site of the cavity may lead to deviations from the model of a perfect decorrelation in XYZ direction. However, the observed decay is exponential, and the decay time shows a linear dependence on the size of the particle (see Figure **3c**), which indicates that any deviations are minor and that a calibration procedure with size standards can account for such geometrical imperfections. Moreover, the nano-cavities are not completely uniform in their depth. Assuming an error of 5% of the cavity depths, this results in an error of 10% in the decay time due to the square dependency of *τ* 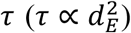. An error of 10% of the decay time may therefore affect the size determination by up to 10%.

In addition, inaccuracies, such surface roughness, can also influence trapping times. However, since the Debye length (<100 nm) and thus the thickness of the electrical double layer is orders of magnitudes larger than the PDMS surface roughness (<5 nm), it can be assumed there is no influence on the trapping time for very small molecules. Furthermore, the electrical double layer itself can have an effect on the effective depth of the cavity and affect the effective excluded volume. However, we do not extract sizes based on absolute cavity depth, which in theory is possible, but we rather use a calibration procedure to account for such effects and to extract sizes. Finally, hydrodynamic coupling to the wall of the cavity can affect the size measurements as well.^30–32^ However, except for the biggest particle tested (50 nm colloids), we are far from a regime where this coupling has a strong effect (see Supporting Information). As such, the effect is smaller than 10% for particles smaller than 20 nm in hydrodynamic radius.

We would like to further note that NDS is robust against measurement noise. Single molecule experiments usually provide data at low signal-to-noise ratios. However, because size information in our approach is extracted from long timescale events, false positive events are exponentially unlikely for longer residence times, as required in our approach. In other words, the likelihood for false particle detection, which mainly happens for the detection of events on short timescales, is minimized, as NDS extracts information from long timescale events. This feature of obtaining data at the high signal-to-noise regime makes NDS robust against measurement noise and thus ensures accurate and reliable measurement of a molecule’s size.

In this work, we have established an approach for the sizing of nano-scale particles using single molecule detection and nanofluidics. The NDS approach harnesses the size-dependent diffusional escape of particles under nano-confinement to obtain size information from the particle’s diffusive properties. Using our theoretical modeling and simulations, we have shown that above a critical timescale, the scaling of the particle’s residence time changes from a power law, which is size independent, to an exponential, size-dependent behavior. This realization forms the basis of our approach and yields a linear behavior of the size of a particle versus its residence time within a nano-cavity. Using a nanofluidic chip combined with confocal microscopy, we have experimentally validated both the exponential scaling behavior for nanoscale particles and biomolecules, and shown that the decay rate follows a linear behavior with respect to the diffusion coefficient. Using such calibration, our approach can yield rapid, accurate, and reliable sizing of particles and biomolecules.

Our NDS approach lines up with other techniques such as FCS and NPT analysis in terms of measurements times, yet no correlation analysis is needed for size determination, and particle sizes from a few nanometers up to tens to hundreds of nanometers can be determined, which is hardly achieved with other techniques. For example, NTA tracks particles only down to ca. 30 nm, while FCS is most sensitive to molecules in the low nanometer regime. By nature, NDS is a single particle counting analysis technique, and such analysis offers the advantage to size heterogenous mixtures with components of different sizes and brightness. In the present paper we have demonstrated how sizes of particles with homogenous size distributions can be measured. However, in future iterations it should be possible to resolve sample mixtures consisting of particles with different sizes. For example, it should be possible to directly resolve different assembly states (e.g., oligomerization states) of a protein based on a difference in their fluorescence intensity. This is afforded by the single molecule sensitivity of confocal detection, which allows for differential thresholding of detected events and creation of particle residence time distributions from subspecies that make up the heterogeneous population. Moreover, the implementation, as demonstrated here, uses fluorescence single molecule detection based on confocal microscopy. However, other readout modalities including total internal reflection microscopy or scattering-based techniques (e.g., iSCAT) can be envisaged as well.

In summary, with NDS we have demonstrated a new single molecule optofluidic approach that allows for a rapid and quantitative sizing of nanoscale objects from a few nanometers up to hundreds of nanometers, opening up potential applications in areas including nanobiotechnology, biophysics, and clinical diagnostics.

## Associated Content

## Supporting Information

The Supporting Information is available free of charge at:

Methods, Experimental setup used for NDS measurements, Simulation code, Mathematical model at short timescales, Mathematical model at long timescales, Considerations on concentration limit and time dependence, Under-sampling at short residence times and data selection criteria, Effect of hydrodynamic coupling within a nano-cavity, Effect of data selection below *t*_c_, Dynamic light scattering (DLS) measurements.

## Author information

### Author Contributions

The manuscript was prepared through the contribution of all co-authors. All authors have given approval to the final version of the manuscript.

## Notes

The authors declare no competing financial interest.

## Acknowledgements

This work was funded by the European Innovation Council under the European Union’s Horizon 2020 Future and Emerging Technologies Open (FET-OPEN) Programme through grant 766972-FET-OPEN-NANOPHLOW (T.P.J.K.). The research leading to these results has further received funding from the European Research Council under the European Union’s Horizon 2020 and the Horizon Europe Framework Programme through Marie Sklodowska-Curie grants MicroSPARK (agreement n°841466; G.K.) and EXO-CHIP (agreement n°101064246; O.V.). G.K. further acknowledges support by the Herchel Smith Funds from the University of Cambridge and the Wolfson College Junior Research Fellowship. This work was also supported by the Engineering and Physical Sciences Research Council (EPSRC) [grant numbers EP/L015889/1] (R.P.B.J.). The authors would also like to thank the EPSRC Centre for Interdisciplinary PhD Training and Research in Nanoscience and Nanotechnology (NanoDTC) for additional funding and the Maxwell Community, University of Cambridge for scientific support.

## Notes

### Competing Interest Statement

The authors have declared no competing interest.

### Summary of Updates

Revised Version.

