## Supporting Information for "Single-molecule sizing through nano-cavity confinement"

### FOR

**Methods**

Simulations

Simulations were performed using a custom written Python script. The diffusion process was modelled as a 1D random walk motion (i.e., unidimensional motion). The simulation tracks the coordinates of the particle as it diffuses in a dead-end channel (i.e., cavity) of depth $d_E$. At each time step $\Delta t$, a new position of the particle is calculated by a Gaussian random number generator with standard deviation $\Delta t \cdot \sigma = \sqrt{2D\Delta t}$, where $D$ is the particle's diffusion coefficient. The entrance of the channel is fixed at position $d_E$. Once the particle passes below the zero position (i.e., bottom wall of the cavity), the position $p$ is updated as $p = |p|$. If the particle's position increases above $d_E$, the path is considered as finished and the time taken to reach the dead end is recorded as the residence time. The probability distribution is estimated from the histogram for a sufficient number of residence times. The long timescale dependence was then fitted to an exponential function to verify the theory. The code is provided in the Section "Simulation code".

Chip preparation for NDS measurements

Fabrication of nano/microfluidic chips was done as detailed in Vanderpoorten et al.[1,2] Briefly, SU-8 photoresist (Type 3025, Micro Resist Technology) was spin coated (Laurell technologies, WS-650) onto a 3-inch silicon wafer (MicroChemicals, Prime CZ-Si). The SU-8 coated wafer was soft baked and treated according to the protocol of the supplier of the photoresist. Microfluidic patterns from a custom-designed film mask (Microlithography) were then projected onto the wafer and the photoresist was exposed for 30 seconds with UV-LED illumination at 365 nm as described in Challa et al.[3] Then the nanofluidic structures of the chip were integrated using the process described in detail by Vanderpoorten al.[1] A custom-built two-photon lithography (2PL) setup was used to write the nanofluidic patterns. After the nanostructures were written, the wafer was baked and developed using standard procedures, and polydimethylsiloxane (PDMS) imprints were created.[4] Briefly, micro-/nanofluidic devices were moulded from the fabricated SU-8 master via soft lithography using PDMS (Sylgard 184; with 10:1 curing agent ratio). After baking, inlets were added using surgical punchers and plasma bonded to coverslip glasses (Menzel coverslips, Grade H1.5). The surface of the coverslips and the PDMS were plasma treated, and afterwards manually pressed on top of each other. PDMS–silica devices were used directly after the plasma bonding step to use their remaining surface hydrophilicity for easier filling of the devices. Before the experiments, the chips were filled by pipetting equal amounts of diluted sample solutions into the inlet areas and equilibrated for 20 minutes. We note that according to previous findings,[5] the PDMS surface roughness can be assumed to be below 5 nm, which should therefore not influence the time spent of particles in the nano-cavity significantly.

Sample preparation for NDS measurements

100 nm and 40 nm fluorescent colloids (FluoSpheres) were purchased from ThermoFisher. α-synuclein oligomers were prepared as described in Vanderpoorten et al.[6] and Hoyer et al.[7] Double-stranded DNA was prepared from two single-stranded DNA oligonucleotides by thermal annealing. Oligonucleotides were synthesized and labelled by Biomers. The sequences were: 5'-GCC TTA **T**TT TCA CTC TTT CCT TTC TTC TTC TCT CTT TTT TTC CCG-3' (top strand) and 5'-CGG GAA AAA AAG AGA GAA GAA GAA AGG AAA GAG TGA AAA TAA 453 GGC-3' (bottom strand); the top strand was labelled with Atto488 at the

thymidine at position 7, shown in bold type. Samples were diluted in deionized water for NDS measurements.

NDS measurements using single-molecule confocal microscopy

NDS measurements were performed using a custom-built single-molecule confocal microscope. A schematic of the setup is shown in **Figure S1**. Briefly, the nanofluidic PDMS–silica device was secured to a motorised microscope stage (Applied Scientific Instrumentation, PZ-2000FT), which was used to position the confocal spot within the chip. The sample was excited using a fibre-coupled and collimated 488-nm wavelength laser (Cobolt 06-MLD, 200 mW diode laser, Cobolt) through a 60X-magnification water-immersion objective (CFI Plan Apochromat WI 60x, NA 1.2, Nikon). The laser intensity at the back aperture of the objective was adjusted to 150 µW. A dichroic mirror (Di03-R488/561, Semrock) was used to separate the excited light from the emission light. Emitted light was collected through the same objective and then passed through a 30 µm pinhole (Thorlabs) to remove any out-of-focus light. The emitted photons were filtered through a band-pass filter (FF01-520/35-25, Semrock) and then focussed onto an avalanche photodiode (APD, SPCM-14, PerkinElmer Optoelectronics) connected to a TimeHarp260 time-correlated single-photon counting unit (PicoQuant). Photon time traces were recorded using the SymPhoTime 64 software package (PicoQuant). Photon recordings were done in T2 mode and the arrival times of photons were measured in respect to the overall measurement start with 16-ps resolution. Data analysis was done using a custom-written Python script. Single-molecule events were identified from the acquired photon stream as fluorescence bursts using a combined maximum inter-photon time ($IPT_{max}$) and minimum total number of photons ($N_{min}$) filter, and an additional Lee filter. The following parameters were used: DNA: $IPT_{max}$ = 0.09 ms, $N_{min}$ = 60, Lee filter = 10; α-synuclein oligomers: $IPT_{max}$ = 0.1 ms, $N_{min}$ = 80, Lee filter = 10; 20 nm colloids: $IPT_{max}$ = 0.2 ms, $N_{min}$ = 100, Lee filter = 15; 50 nm colloids: $IPT_{max}$ = 0.2 ms, $N_{min}$ = 150, Lee filter = 15. Photon counts were averaged into 1-ms binned time traces.

**Experimental setup used for NDS measurements**

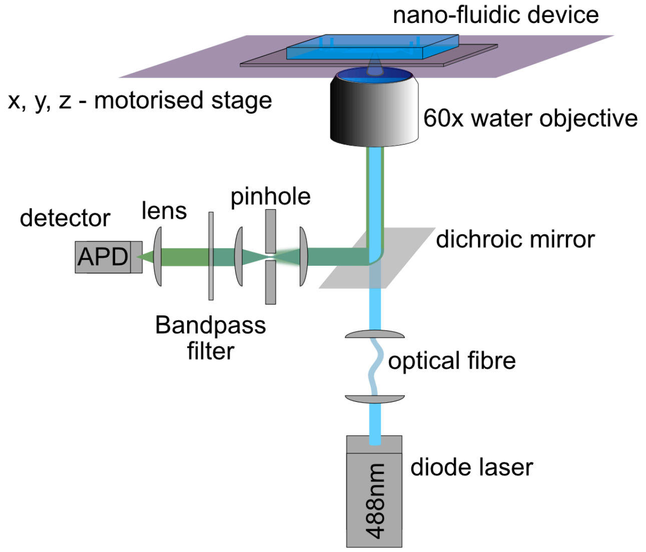


**Figure S1.** Schematic of the confocal system used for NDS measurements. A 488-nm diode laser is used to excite the sample through an objective, and a single-photon counting avalanche photodiode (APD) is used to register emitted fluorescence photons. The confocal detection volume of the microscope is positioned within the nano-cavities of the chip with the aid of a motorised stage. Details of the setup are described in the Methods section.

#### Simulation code

```
# -*- coding: utf-8 -*-
"""
Created on Tue Apr 14 10:43:38 2020

@author: Quentin.Peter
"""

import numpy as np
import scipy.ndimage.measurements as msr
import matplotlib.pyplot as plt
#%%
m = [0,0,0,0,0,0,0]
#nbr_runs = [100,1000,10000,50000,100000,500000,1000000]
nbr_runs = [100000]
m = np.zeros(np.shape(nbr_runs)[0])
ii = 0
N = 100
for number_runs in nbr_runs:
    wall = np.sqrt(100)
    all_counts = np.zeros(2, dtype=int)
    #number_runs = 100000

    while all_counts[1] < number_runs / 2:
        traces = np.random.normal(size=(2**27)) #1GB
        # Create traces from steps
        traces =  np.cumsum(traces, -1)

        # Boundary condition (keep traces between -wall and wall)
        traces = np.abs(traces + wall)
        invert = (traces // (2 * wall)) % 2 == 0
        traces -= (traces // (2 * wall)) * (2 * wall)
        traces[invert] = 2 * wall - traces[invert]
        del invert
        traces -= wall

        # label traces
        for i in range(2):

            if i==0:
                lbl, _ = msr.label(traces > 0)
            else:
                lbl, _ = msr.label(traces < 0)

            # remove last trace
            if lbl[-1] !=0:
                lbl[lbl == lbl[-1]] = 0

            indices = np.ravel(np.argwhere(np.bincount(lbl) > N))[1:]
            # Compute length
            lengths = np.bincount(lbl)[1:]
            length_count = np.bincount(lengths)
            # Merge counts
            if len(length_count) > len(all_counts):
                all_counts, length_count = length_count, all_counts
            all_counts[:len(length_count)] += length_count
            # Free memory
            del lbl, lengths, length_count
```

```
        idx_transition = np.ravel(np.argwhere(np.diff(np.asarray(traces > 0, int)) > 0))
        idx_transition = idx_transition[indices - 2] + N + 1
        plt.hist(traces[idx_transition], 50)
        del traces
        print(2 * all_counts[1])
    # Compute log bins
    nbins = 30
    basis = np.exp(np.log(len(all_counts))/nbins)
    X = np.arange(len(all_counts))
    lbl = np.asarray(np.log(X) / np.log(basis), int)
    bins = msr.minimum(X[1:], lbl[1:], index = np.unique(lbl[1:]))
    mid_bins = msr.mean(X[1:], lbl[1:], index = np.unique(lbl[1:]))
    measure = msr.mean(all_counts[1:], lbl[1:], index = np.unique(lbl[1:]))
    # fit exponential and power law
    fit = np.polyfit(np.log(mid_bins[mid_bins < wall**2]), np.log(measure[mid_bins < wall**2]), 1)
    fit2 = np.polyfit(mid_bins[mid_bins > wall**2], np.log(measure[mid_bins > wall**2]), 1)
    m[ii] = fit2[0]
    ii += 1

plt.plot(nbr_runs, np.abs(m))
```

**Mathematical model at short timescales**

In the case of a free random walk without constraints, the system is scale invariant for the first return to the origin, as explained in the first chapter of "A guide to first-passage processes" by Redner.[8] This scenario corresponds to a special case of the first-time-hitting model in which $\Delta y = 0$. If $\Delta y$ is small, but not equal to zero, the system shows a scale dependence on *D*. However, for a fixed minimum detection time *T* required to detect the particle, the system becomes scale invariant again. This is because, within the minimum detection time *T*, the particle has penetrated the cavity by $\Delta y$, and this mean penetration distance itself depends on $\sqrt{2DT}$. Hence, the system becomes scale invariant again according to:

$$p(t, D) = \frac{\Delta y(D)}{\sqrt{4\pi D t^3}} \propto \frac{\sqrt{2DT}}{\sqrt{4D\pi t^3}} \propto t^{-\frac{3}{2}} \tag{S1}$$

Another formal approach, which describes the power law dependency of the asymptotic behaviour at $\Delta y = 0$, to derive this behaviour can be found in the first chapter of Redner as well.[8] The dependency of having a particle passing through a point at a distance $\Delta y$ in a time interval $\Delta t$ is given by $P(\Delta y, \Delta t)$. For the case of $\Delta y = 0$, it can be described as:

$$P(0, \Delta t) \propto \int_0^{\Delta t} (4\pi D t)^{-\frac{d}{2}} dt \sim \begin{cases} A_d \Delta t^{\left(1-\frac{d}{2}\right)}, & d < 2 \\ A_2 \ln(\Delta t), & d = 2 \end{cases} \tag{S2}$$

where $d$ is the dimension of the system, and $A_d$ is of the order of 1 and does not play a role in the asymptotic behaviour. The survival probability ($S(\Delta t)$) is given by:

$$S(\Delta t) \sim \begin{cases} \frac{1}{A_d \Delta t^{\left(1-\frac{d}{2}\right)}}, & d < 2 \\ \frac{1}{A_2 \ln{(\Delta t)}}, & d = 2 \end{cases} \tag{S3}$$

The first passage probability or density at the origin ($p(t)$) is then formulated using the following relationship, $1 - S(\Delta t) \sim \int^{\Delta t} p(t)dt$, which leads to:

$$p(t) = -\frac{\partial S(t)}{\partial t} \propto \begin{cases} t^{\frac{d}{2}-2}, & d < 2 \\ \frac{1}{t \ln^2 t}, & d = 2 \end{cases} \tag{S4}$$

Accordingly, $p(t)$) is proportional to $t^{-\frac{3}{2}}$ and, hence, scale invariant.

**Mathematical model at long timescales**

The problem of modelling diffusion within a confined geometry (i.e., nanocavity) is equivalent to the problem of modelling diffusion of particles in a straight channel which has two exits. This is due to boundary reflexion conditions at the end of the cavity. Such a scenario has been previously described by Balluffi et al.[9] (Eq. 5.47 therein). The size of this system is equivalent to our system with reflective boundary conditions if we take into account $2d_E = L$. Accordingly, the position *x*- and time *t*-dependent concentration *C* within the channel cross-section can be written as:

$$C(x,t) = \frac{4C_0}{\pi} \sum_{j=0}^{\infty} \sin\left(\frac{(2j+1)\pi x}{L}\right) \exp\left(-\frac{(2j+1)^2\pi^2}{L^2} Dt\right) \tag{S5}$$

with $C_0$ being the concentration at time point and position $t=0$, $x = 0$.
For long timescales, such that $\frac{\pi^2}{L^2} Dt \gg 1$, other contributions than $j = 0$ can be neglected. The concentration is then:

$$C(x,t) = \frac{4C_0}{\pi} \sin\left(\frac{\pi x}{L}\right) \exp\left(-\frac{\pi^2}{L^2} Dt\right) \tag{S6}$$

We can integrate the system over the length of the channel *L* to obtain $C_{tot}$:

$$C_{tot}(t) = \int_0^L C(x,t) = \frac{4C_0}{\pi^2} 2L \exp\left(-\frac{\pi^2}{L^2} Dt\right) \tag{S7}$$

Finally, the change of $C_{tot}$ over time is:

$$\frac{dC_{tot}(t)}{dt} = \frac{4C_0}{\pi^2} 2L \left(-\frac{\pi^2}{L^2} D\right) \exp\left(-\frac{\pi^2}{L^2} Dt\right) \propto \exp\left(-\frac{t}{\tau}\right) \tag{S8}$$

The decay time is given by $\tau$, and depends only on the size *L* and the diffusion coefficient *D*:

$$\tau = \frac{(2d_E)^2}{\pi^2 D} \tag{S9}$$

For long timescales ($t \gg 2d_E^2/\pi^2 D$), the escape time decreases exponentially. The decay time $\tau$ is then inversely proportional to the diffusion coefficient *D*.

An interesting, alternative derivation of the theory is presented by Redner et al.[8] The first passage time is developed within a similar restricted geometry and, despite a slightly different approach, the same result is achieved for long-timescale limits with a decay time $\tau$ of $\frac{(2d_E)^2}{\pi^2 D}$.

**Considerations on the concentration limit and time dependence**

The concentration limits for the sizing of single particles, both at high and low concentrations, depends on the probability for a particle to be within the nano-cavity. At high concentrations, if more than one particle is present within the cavity at a time, the observed dwell time distribution will be skewed towards longer residence times. Hence, this will skew the residence time probability distribution. To determine what is an acceptable concentration, we assume arbitrarily that if 10% of the detected events are made up by two particles or more, the error becomes too large to characterize the size accurately. This is defined as the high concentration limit. As represented in **Figure S2a**, the high concentration limit for our system is on the order of 5.5 μM. The probability distribution shown in the figure is based on the cumulative distribution function derived from Poisson statistics[10] at an event rate of $k = 1$:

$$f(k) = \exp(-\mu)\frac{\mu^k}{k!} \tag{S10}$$

where $\mu$ is the mean particle concentration in the cavity.

In the low concentration regime, time becomes an important factor for the sizing of particles. **Figure S2b** represents the probability to detect a particle at a given concentration. Hence, for a given time, the probability to detect enough events to calculate the size of our particle decreases with concentration. **Figure S2c** shows the probability to detect particles assuming a diffusion coefficient *D* for a standard protein such as bovine serum albumin (BSA, $D = 5.9 \cdot 10^{--7}$ $cm^2/s$). Assuming at least 1000 events are necessary to be able to size a particle in less than 30 min, the concentration range needs to be on the order of nM or higher.

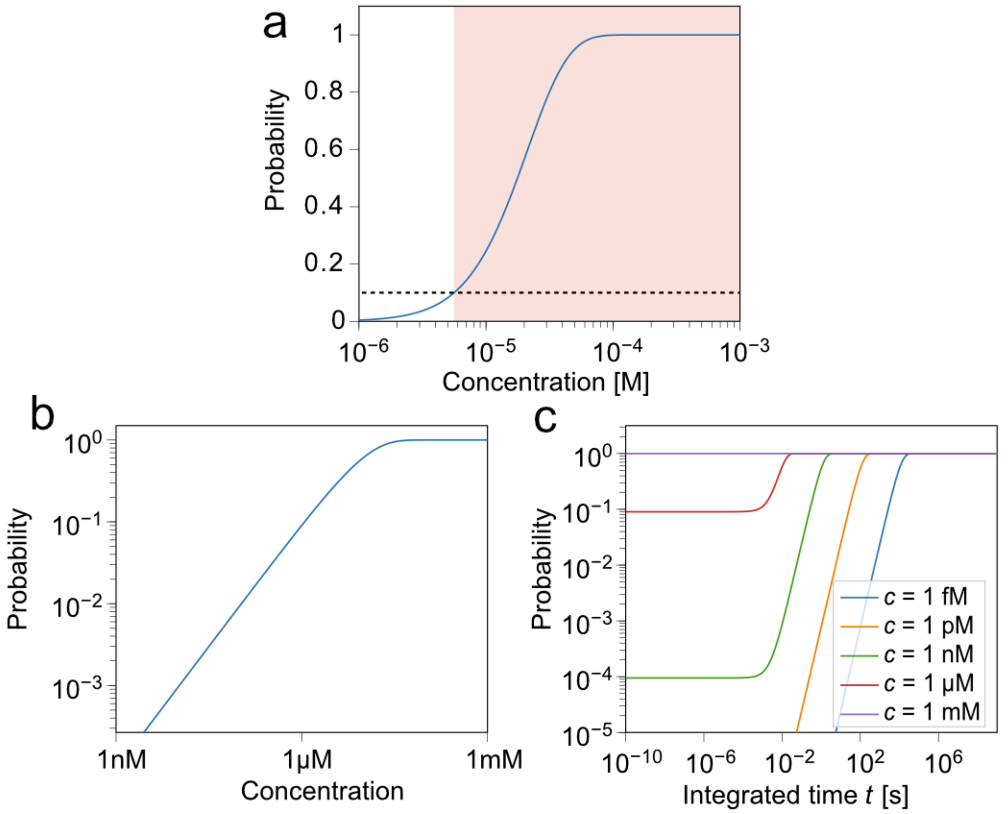


**Figure S2. (a)** Evolution of the Poisson probability distribution to encounter two particles inside a nano-cavity at the same time. The dimensions of the cavity were 600 nm x 750 nm x 350 nm. The red area denotes the regime where more than 10% of events have a likelihood to consist of two or more molecules. The dashed line represents the 10% probability threshold. **(b)** Probability to encounter at least one particle as a function of particle concentration. **(c)** Evolution of the probability distribution function to encounter at least one particle during an integration time $t$ for various particle concentrations (as denoted). The particle was assumed to have a diffusion coefficient of $D = 5.9 \cdot 10^{-7}$ cm$^2$/s.

#### Under-sampling at short residence times and data selection criteria

Due to the nature of the measurement, short residence time events are under-sampled. This is because the probability for an event to be above a certain noise threshold level increases with the time the particle remains in the confocal detection volume. Hence, shorter events are under-sampled in comparison to longer events. This is evident in the peak-shaped distribution of residence times when histogramming detected residence times in our experiments. **Figure S3** illustrates this behaviour. The particle residence time distributions all exhibit a short timescale flank, which represents the under-sampled regime, then peak, and then follow an exponential decay. In our data analysis, we have discarded the under-sampled region and fitted the decay from the peak of the shape until the high end of residence times. An additional filter criterion was used, which involved the removal of statistically irrelevant events (i.e., residence time intervals, after falling below 5 counts in the histogram, were not taken into account in the analysis).

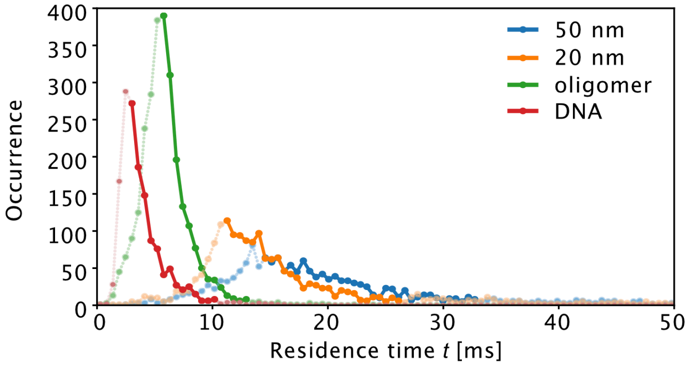


**Figure S3.** Occurrence of detected single particle residence times for 50 nm colloids, 20 nm colloids, α-synuclein oligomers, and DNA oligos. The regions that were used to fit the exponential decay are shown as solid lines. The discarded points from the short timescale flank and those that were statistically irrelevant are shown in light colour and as dotted lines.

#### Effect of hydrodynamic coupling within a nano-cavity

For nanoparticles diffusing in nano-cavities, which exhibit similar dimensions as the nanoparticle of interest, hydrodynamic coupling effects can occur, especially close to the cavity walls. These can affect the diffusive behaviour of the particles. Faxén and Brenner have described this phenomenon.[11–13] Accordingly, the diffusion coefficient $D$ is affected by:

$$D_{\parallel}(h) = D[1 - \frac{9}{16}\frac{R}{h} + O(\frac{R^3}{h^3})] \qquad \text{(S11)}$$

where $D_{\parallel}$ is the diffusion coefficient parallel to the wall and $h$ is the distance of the particle to the side of the wall, and $R$ is the radius of the particle. The average diffusion coefficient is then obtained by calculating the integral over all distances of the particle to the wall $h$. Hence, for a nano-cavity with a width $w$, the average diffusion coefficient $\widetilde{D}$ can be calculated as follows:

$$\widetilde{D} = \frac{2}{2(\frac{w}{2} - R)}\int_R^{w/2} D_{\parallel}(h)\,dh \qquad \text{(S12)}$$

$$\widetilde{D} = a(R,w)\,D = \frac{1}{(\frac{w}{2} - R)}\left[\frac{w}{2} - \frac{9}{16}R\ln\frac{w}{2} - R + \frac{9}{16}R\ln R\right]D \qquad \text{(S13)}$$

with $a(R,w) = \widetilde{D}/_{D}$ describing the change in average diffusion coefficient $\widetilde{D}$ with respect to $D$.

**Figure S4** depicts the change in $a(R)$ as a function of particle radius $R$ for a fixed well width of $w$ = 700 nm. Accordingly, hydrodynamic effects reduce the observed $\widetilde{D}$ by about 2% for 1 nm particles ($a$ = 0.98) and up to 20% for 50 nm particles ($a$ = 0.8).

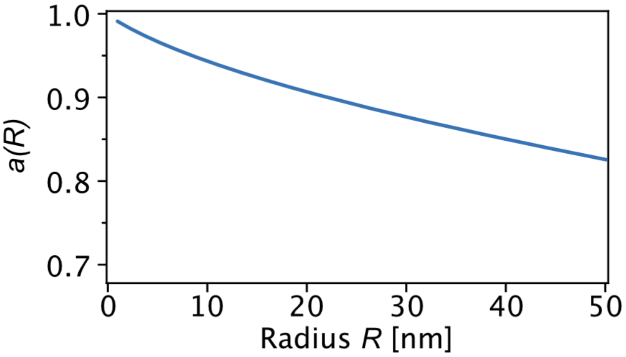


**Figure S4.** Change in $a(R)$ as as a function of particle radius $R$ for a well width of $w = 700$ nm.

#### Effect of data selection below $t_c$

For measuring particle sizes in experiments, the critical time $t_c$ is not known, as the particle size is typically not known. Hence, in the experiments shown in **Figure 3** (main text), $t_c$ was not considered when calculating the decay time. However, due to under-sampling effects at short time scales (see **Figure S2**), the points in the short times scale regime close to $t_c$ and below are already mostly discarded. Hence, in most cases, the cut-off at or around $t_c$ is already introduced by the data selection procedure (see above). Nonetheless, it is possible to estimate $t_c$ from the data by an iterative approach. This is done by first performing a calculation of the size of the particle of interest including all data points, and using a calibration procedure, to calculate $t_c$. In a second step, $t_c$ is applied to the data set to remove data points below $t_c$, and the size of the particle is calculated from the subset of points. **Figure S5** depicts the effect of removing data points below $t_c$. In our data set in **Figure 3**, only the 50 nm colloids and the DNA sample exhibited data points below $t_c$ (5 data points for the 50 nm colloid and 1 data point for the DNA). For these samples, the calculated change in size, by considering $t_c$, was found to be smaller than 1.5% and 0.52%, respectively.

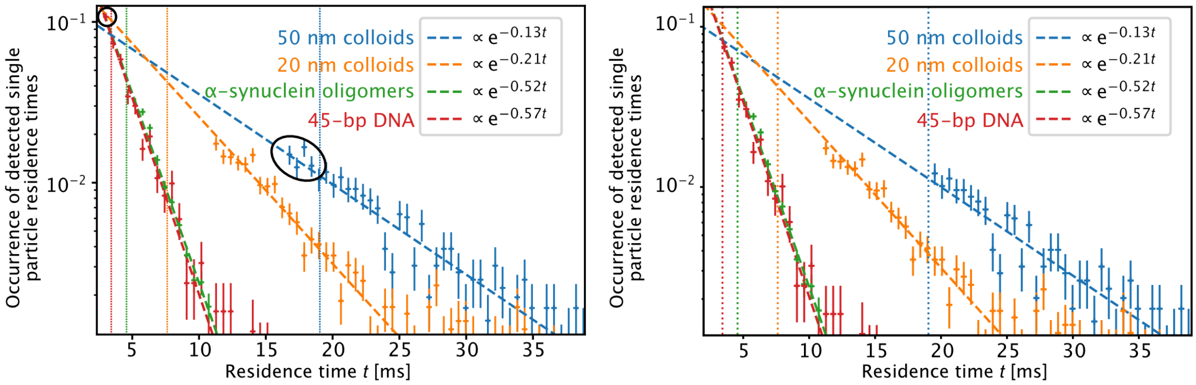


**Figure S5.** Effect of removing data points below the critical time $t_c$. **(a)** Residence time decay histograms taking into account all data points (as shown in Figure 3c in the main text). **(b)** Residence time decay histogram where all data points below $t_c$ were removed. Only the 50 nm colloid and the DNA sample exhibit points below $t_c$ (highlighted in panel a). Removal of these data points only marginally affected the slopes of the exponential fits (see insets).

#### Dynamic light scattering (DLS) measurements

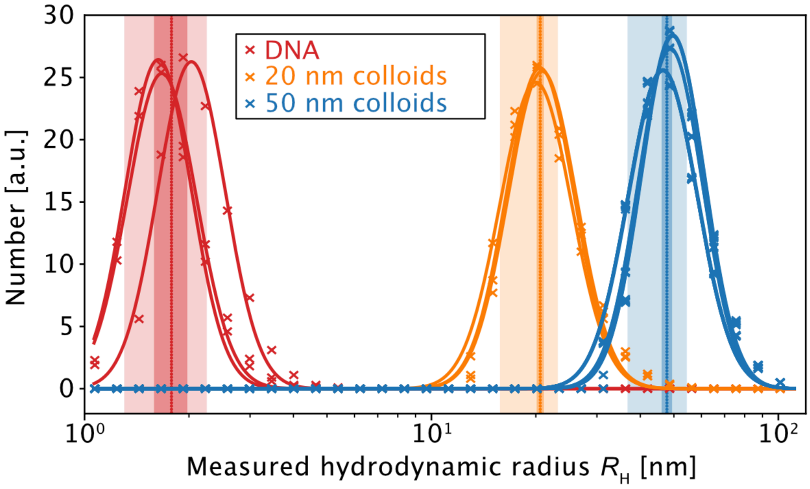


**Figure S6.** Hydrodynamic radii $R_H$ for DNA, 20 nm colloids and 50 nm colloids as measured by DLS. Measurements were done in triplicates. Mean values of hydrodynamic radii are depicted as vertical lines (DNA: 1.9 nm, 20 nm colloids: 20.6 nm, and 50 nm colloids: 47.7nm). Error bars are shown as shaded areas. The dark-shaded error bars correspond to the standard deviation of triplicate measurements. The light-shaded error bar regions correspond to the standard deviation of a Gaussian fit of the distribution in log-space. See also Table 1 (main text).
